# Overcoming confounding plate effects in differential expression analyses of single-cell RNA-seq data

**DOI:** 10.1101/073973

**Authors:** Aaron T. L. Lun, John C. Marioni

## Abstract

An increasing number of studies are using single-cell RNA-sequencing (scRNA-seq) to characterize the gene expression profiles of individual cells. One common analysis applied to scRNA-seq data involves detecting differentially expressed (DE) genes between cells in different biological groups. However, many experiments are designed such that the cells to be compared are processed in separate plates or chips, meaning that the groupings are confounded with systematic plate effects. This confounding aspect is frequently ignored in DE analyses of scRNA-seq data. In this article, we demonstrate that failing to consider plate effects in the statistical model results in loss of type I error control. A solution is proposed whereby counts are summed from all cells in each plate and the count sums for all plates are used in the DE analysis. This restores type I error control in the presence of plate effects without compromising detection power in simulated data. Summation is also robust to varying numbers and library sizes of cells on each plate. Similar results are observed in DE analyses of real data where the use of count sums instead of single-cell counts improves specificity and the ranking of relevant genes. This suggests that summation can assist in maintaining statistical rigour in DE analyses of scRNA-seq data with plate effects.

## Introduction

Single-cell RNA sequencing (scRNA-seq) is increasingly being used to study molecular biology at the cellular level. RNA is isolated from individual cells and reverse-transcribed into cDNA fragments that are sequenced using massively parallel technologies [22]. Mapping of the sequence reads to a reference genome allows quantification of gene expression in each cell based on the number of reads assigned to each gene. The count data can then be analyzed to identify cell subtypes by clustering of the gene expression profiles; to identify highly variable genes contributing to cell-to-cell heterogeneity; and to identify differentially expressed (DE) genes between groups of cells. The ability to assay expression profiles for individual cells provides scRNA-seq studies with biological resolution that cannot be matched by bulk RNA-seq experiments on cell populations. However, this comes at the cost of high technical noise due to difficulties in sequencing low input quantities of RNA from single cells.

Most existing scRNA-seq protocols are based either on microwell plates [17] or microfluidics systems like the Fluidigm C1 [18]. In the former, cells are placed into individual wells, typically with automated approaches such as fluorescently activated cell sorting (FACS). The entire plate is processed to generate sequencing libraries for all cells at once. In the latter, individual cells are captured in separate reaction chambers within the C1 chip (for simplicity, each chip will be treated as being equivalent to a plate in the following text). All cells on each chip are then simultaneously processed into libraries. Each plate or chip can only handle a limited number of cells, so data are usually generated for multiple plates across several biological groups. All cells on each plate usually come from the same population (e.g., cell culture or animal) from a single biological group. If replicate populations are present in a group, a separate plate is typically used to process the cells from each population.

The experimental design described above is motivated largely by practicality. It is logistically simpler to track and process cells when each plate corresponds to one population of one group, compared to partitioning the plate into different groups. For example, FACS is typically performed such that all cells on a plate are obtained by gating on a single subpopulation. In the C1, cells are captured randomly in reaction chambers such that the identity of the cell in each chamber cannot be pre-specified. Ambiguity in determining the group for each cell is avoided by only using cells from the same group on each chip. However, this design can result in a “plate effect” where uncontrollable experimental variables have a consistent effect on the observed counts for all cells on each plate but a variable effect across different plates. These variables can be technical in origin, such as differences in library preparation or sequencing between plates; or biological, due to inherent variability in gene expression across replicate populations on different plates.

One common analysis applied to scRNA-seq data involves identifying DE genes between groups of cells. This does not strictly require single-cell resolution, and is more conventionally performed with bulk RNA-seq data. Nonetheless, there are some benefits to using single-cell data in this context. Rare cells are more easily sequenced with single-cell protocols, and irrelevant cells from contaminating populations can be screened out before analysis. It is also more practical to use existing scRNA-seq data compared to generating bulk data exclusively for a DE analysis. The analysis itself is typically performed using methods like edgeR [19] and DESeq2 [12], which were originally designed for bulk data; or with methods such as Monocle [23] and SCDE [6], which were designed explicitly for single-cell data. Putative DE genes are defined as those that exhibit significant differences in the average counts between two or more groups of interest.

The presence of plate effects complicates the parametrization of the experimental design in the DE analysis. Obviously, such effects are undesirable as they arise via uncontrolled experimental variability. This increases the estimated variance and reduces the power to detect DE genes. However, plate effects cannot be easily removed during statistical modelling since they are confounded with the groups of interest [4, 24]. In the most extreme case, consider a data set with multiple groups where each group is comprised of cells from a single plate. Changes in gene expression due to plate effects will be indistinguishable from genuine DE between groups. Any attempt to remove the former will compromise the detection of the latter. This problem is still present in data sets with a few plates (e.g., 2-3) in each group, where stochastic plate-to-plate variability may explain some or all of the apparent DE between groups. Many studies ignore the plate effects and treat cells directly as replicates in the DE analysis, such that the variance is estimated across cells in each group [8–10]. This strategy is not statistically rigorous as the variability due to systematic differences between plates will not be modelled properly.

This article demonstrates that plate effects must be handled appropriately to maintain the statistical validity of a DE analysis. In a simple simulation, all analyses that ignored the plate effects failed to control the type I error rate. This was caused by dependencies between cells on the same plate which compromised inferences from the fitted model. To avoid detection of spurious DE, a summation approach was proposed whereby counts for all cells on each plate were summed prior to the DE analysis. This restored control of type I error across a range of simulation scenarios without compromising DE detection power. Similar effects were observed in real data where summation improved detection specificity and the ranking of relevant genes.

## Plate effects result in false positives in a DE analysis

### Description of the experimental design and analysis strategies

Consider an experimental design consisting of multiple biological groups of interest. Each group consists of cells sorted onto multiple plates, where all cells on each plate come from an independent population of a single group. As previously mentioned, this is not an uncommon setup due to the logistics of the experimental protocol, e.g., cell sorting onto a plate with FACS, cell capture with the C1. Further assume that a plate effect exists in the data set, whereby the expression of each gene in all cells of a given plate is modified in a gene- and plate-specific manner. The use of gene-specific effects is motivated by the presence of biological variability between replicate populations, where uncontrolled differences in cell culturing or animal treatment can affect expression in a gene-dependent manner.

Let *Y*_*ijkg*_ be a random variable representing the read count for gene *i* in cell *j* in plate *k* of group *g*. Define *δ*_*ikg*_as a random variable representing the gene-/plate-specific effect for gene *i* in plate *k* of group *g*. Assume that *δ*_*ikg*_ for each plate is independently sampled from a distribution of positive values with a mean of unity and non-zero variance. This is reasonable as each plate contains cells from an independent population and is processed independently. Further define *θ*_*jkg*_ as a random variable representing the bias in cell *j* in plate *k* of group *g*, also sampled from a distribution of positive values with a mean of unity and non-zero variance. This is independent of *δ*_*ikg*_ as it represents the cell-specific bias (e.g., library size, composition bias) within each plate, which is separate from the gene-specific plate effects. *Y*_*ijkg*_ is sampled independently for each gene in each cell, conditional on the observed values of *δ*_*ikg*_ and *θ*_*jkg*_. In the following text, we refer to var(*Y*_*ijkg*_|*δ*_*ikg*_, *θ*_*jkg*_) as the conditional variance, while the conditional expectation is defined as

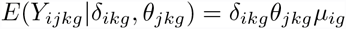

where *µ*_*ig*_ as the expected read count for gene *i* in group *g*. This represents the impact of plate effects on the data. For each gene, the mean for all cells on a given plate is scaled by the same value, *δ*_*ikg*_, which introduces systematic differences between plates. Such differences cannot be removed by scaling normalization, e.g., based on library size or with more sophisticated methods [1,20]. This is because these normalization methods compute a single scaling factor for each cell to adjust for the cell-specific *θ*_*jkg*_. In contrast, the plate effect varies across genes and cannot be removed by a single factor for each cell.

One common aim of the data analysis for this experiment is to identify DE genes between the biological groups of interest. However, this is complicated by the presence of confounding plate effects. Consider an experimental design with one plate from each of two groups. A fold-increase in *δ*_*ikg*_ between the two plates cannot be distinguished from a matching increase in *µ*_*ig*_ between the two groups. Any attempt to remove the former will affect detection of genuine DE in the latter. Even with multiple plates per group, this problem is still present as stochastic increases in *δ*_*ikg*_ for all plates in a group can explain part or all of the observed DE between groups.

A common strategy is to ignore the plate effects and treat cells from different plates in the same group as replicates. This is most obviously applied in a one-way layout with clearly defined groups. It is also relevant to multi-factor designs with more plates than model coefficients, as these designs will contain some level of replication between plates and thus between their constituent cells. DE genes can then be detected using software designed for bulk data (e.g., DESeq) or with dedicated single-cell methods (e.g., Monocle). Alternatively, a more sophisticated but lesser-used approach involves fitting a mixed model to the counts for each gene [24]. The plate of origin is treated as a random effect while the groups are treated as fixed effects. This accounts for plate-to-plate variability while still allowing detection of DE between groups. However, the performance of these strategies has yet to be rigorously assessed on scRNA-seq data sets that contain plate effects.

### Testing DE analysis methods on simulated scRNA-seq data

We designed simulations to mimic the characteristics of a real scRNA-seq data set [8]. We generated counts for 50-100 cells in each plate, for an experiment containing three plates in each of two groups. Counts for each cell were conditionally NB-distributed and the plate effect was log-normally distributed. Parameters of both distributions were estimated from real data – see Supplementary Figure S1, Section 1 of the Supplementary Materials for details.

DE analyses of the simulated data were then performed using edgeR, DESeq2 [12], voom [11], Monocle, MAST [3] and generalized linear mixed models (GLMMs) from the lme4 package [2]. edgeR, DESeq2 and voom were originally designed for analyzing bulk RNA-seq data. edgeR and DESeq2 fit NB generalized linear models (GLMs) to the counts for each gene, while voom uses a normal distribution to model log-counts per million (log-CPMs) with precision weights. Monocle and MAST were explicitly designed for analyzing scRNA-seq data, and operate on pre-processed expression values such as (log-)CPMs. (In particular, Monocle was designed to detect DE across a pseudo-temporal ordering of single cells, e.g., during differentiation, but the same statistical framework can be applied to arbitrary covariates.) The GLMM approach is more general, and uses NB distributions to model the counts with random plate effects and fixed group effects.

Each method was applied to the simulated data to test for DE between groups. For all methods, the experimental design was parameterized as a one-way layout with two groups. All cells within each group were treated as direct replicates – the plate of origin within each group was ignored. edgeR was run twice, using either the quasi-likelihood (QL) framework [14] with robust empirical Bayes (EB) shrinkage [16] of the QL dispersions, or the likelihood ratio test (LRT) [15] without any EB shrinkage of the NB dispersions. Similarly, the analysis with voom was repeated after estimating the correlation between cells on the same plate [21]. The implementation details for each method are described in Section 2 of the Supplementary Materials.

In this simulation, the null hypothesis is true for each gene *i* as *µ*_*ig*_ is constant for all *g*. Any rejections of the null represent type I errors, i.e., false positives. For a specified type I error rate *α* = 0.01, the observed error rate was defined as the proportion of all genes with a *p*-value below *α*. This was averaged across 10 simulation iterations to obtain a stable estimate of the observed type I error rate for each method. A method was considered to be liberal if its observed error rate in the simulation was above the specified *α*. Note that this evaluation is only possible for methods that compute *p*-values for each gene – Bayesian methods [6, 25] are not directly comparable and so are not tested here.

### Type I error control is lost by all methods

The observed type I error rate exceeds the specified threshold in all methods (Figure 1). For most methods, the discrepancy between the observed and specified rates is greater than an order of magnitude. This liberalness is attributable to the plate effect rather than any inherent fault with the methods. When the simulations are repeated without any plate effect, the observed rates for all methods are substantially closer to the specified level, if not below it. These results suggest that DE analyses will perform poorly if the plate effect is simply ignored. Loss of type I error control will result in an unacceptable number of false positives in the final set of DE genes.

**Figure 1.**
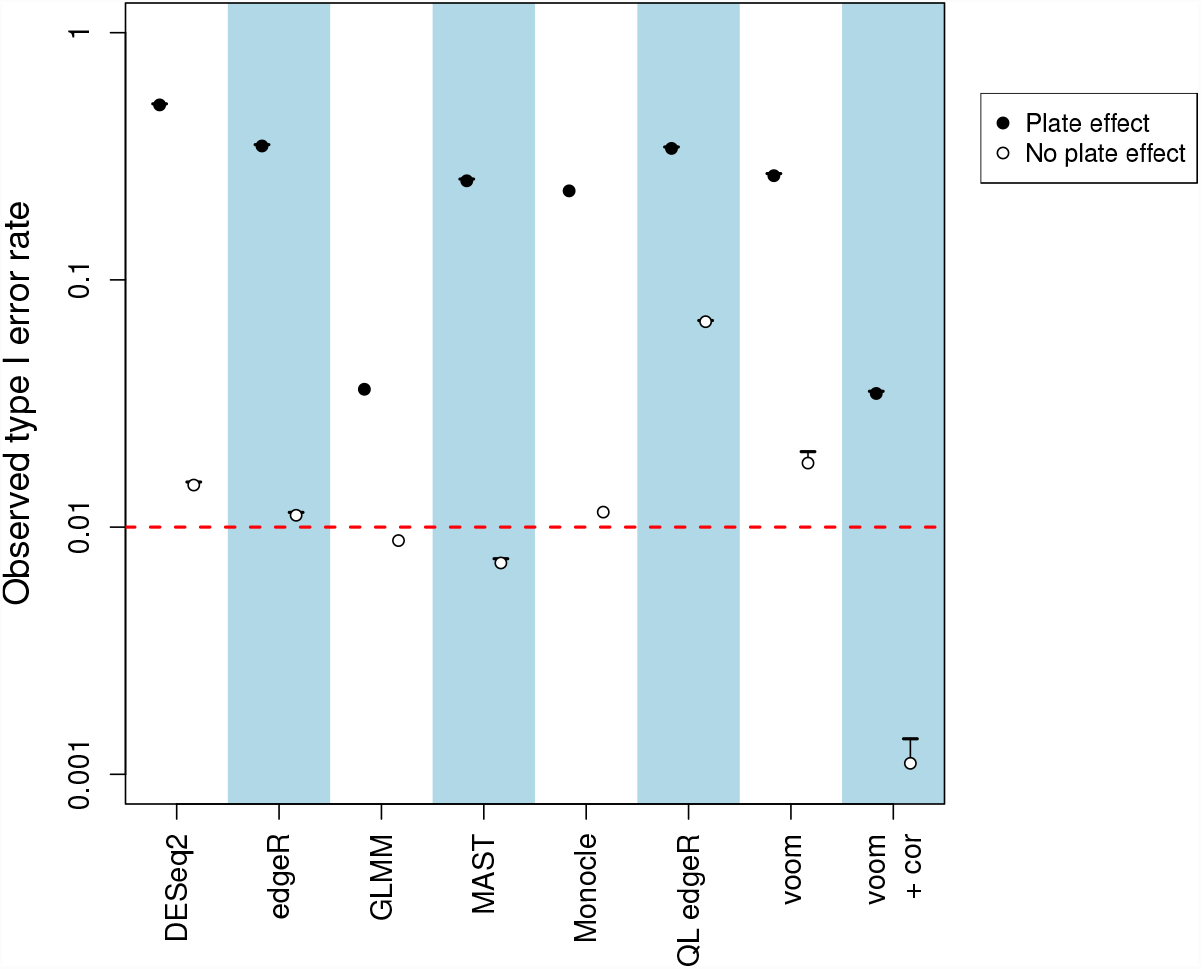
Observed type I error rates for each method on simulated data, in the presence or absence of a plate effect. Error rates are shown on a log scale and represent the average across 10 simulation iterations. Each error bar represents the standard error of the log-average rate. The threshold of 0.01 is represented by the red line. Only one iteration was used for Monocle and GLMMs due to their long run times.

Loss of error control is caused by plate-induced dependencies in the statistical model. In the one-way layout, the count of each gene in each cell is modelled by a distribution with a mean equal to the product of a gene- and group-specific expression term (the estimate of *µ*_*ig*_) and a cell-specific scaling factor (*θ*_*jkg*_). Most DE analysis methods assume that, for any given gene, the counts for all cells are independently sampled from these distributions. However, this is clearly not the case when a plate effect is present. For each gene, the true mean of the sampling distribution for each cell is scaled by a plate-specific factor *δ*_*ikg*_, such that the residuals of the fitted model for cells on the same plate are more similar than expected under independence. (Conditional independence between cells is irrelevant here, as the plate of origin is not used in the model parametrization.) This reduces the amount of information in the data set as cells on the same plate are effectively redundant. Methods that assume independence will overstate the information available to estimate model parameters such as the group-specific means, the variance or the NB dispersion. This results in overconfident inferences and liberalness during hypothesis testing.

The problem can also be described in terms of the residual degrees of freedom (d.f.) that are available in the fitted model. The simulation uses an experimental design with 6 × 50 = 300 cells and two coefficients in a one-way layout. If all cells were truly independent – as is assumed by most methods – this design would provide 298 residual d.f. At the other extreme, consider the scenario where all cells on a plate have identical counts, i.e., there is no noise or heterogeneity between cells. This means that there are only six independent samples (one cell per plate, as all other cells on that plate are redundant) and only four residual d.f. This is substantially smaller than the expected 298 d.f., meaning that the uncertainty of various parameter estimates will be underestimated. For example, the use of edgeR with the LRT assumes that sufficient residual d.f. are present to estimate the dispersion for each gene separately, without requiring EB shrinkage to share information between genes. This is no longer justifiable if only four d.f. are available.

The GLMM approach and voom with correlations exhibit the best performance of all tested methods in the presence of a plate effect (Figure 1). This is because they explicitly account for the dependencies between cells on the same plate during variance estimation. Nonetheless, both methods are still liberal, with observed type I error rates that are approximately three-fold higher than the specified threshold. This is attributable to difficulties in estimating the variance of the random effect (for the GLMM approach) or the correlation (for voom) in data sets with small numbers of replicate plates. Underestimation will result in loss of type I error control as the dependencies will not be fully modelled. For voom, the effect of imprecise estimation with few plates is mitigated by sharing information across genes to obtain a consensus value of the correlation – however, this leads to biases when the true correlation varies between genes. GLMMs are also prohibitively slow to fit, taking several hours to run for a single simulation iteration.

We also tested the performance of each method with other simulation settings. To this end, we repeated the above simulation after halving the magnitude of the plate effect; increasing the variability of library sizes across cells; increasing the variability in the number of cells per plate; increasing the number of plates in each group; or sampling counts from a zero-inflated negative binomial distribution, with parameters estimated from real data [26]. In all scenarios, type I error control was not maintained by any method (Supplementary Figure S2). This suggests that our results are generally applicable to different scRNA-seq data sets.

## Improved performance with summation across cells in each plate

### Error control can be restored by summing over cells

A simple solution presents itself for restoring error control in the presence of plate effects. Firstly, the counts for each gene are summed across all cells within each plate. These count sums are then used directly in the DE analysis, where the plates themselves are treated as replicate samples for each biological group. The use of plate-based observations avoids dependencies between samples in the statistical model. This is because the plate effect is independently sampled for each plate and will not introduce unexpected similarities between count sums for different plates. Similarly, the counts for cells within each plate are conditionally independent and will not provide any information on the counts in other plates. Independence of the count sums fulfills the expectations of the analysis methods and ensures that the number of residual d.f. is not overestimated. Summation has the additional benefit of increasing the size and precision of the counts. This makes the data more amenable for analysis with existing methods designed for bulk data.

Summation substantially reduces the liberalness of the DE analysis methods in the simulation (Figure 2). Overconfident estimation of model parameters is avoided due to the presence of independent count sums. Similar results are observed in the alternative simulation scenarios (Supplementary Figure S3). Note that some mild liberalness is still observed for edgeR and DESeq2 – this is because the count sums are not NB-distributed which results in some inaccuracy during modelling. In contrast, type I error control is fully restored for voom as it is more accurate for large counts and log-normally-distributed plate effects. The other methods are not used here, for various reasons – voom with correlations and GLMMs cannot be applied on count sums from independent plates, as the plate-level blocking factor would be confounded with the random error; Monocle and MAST are designed for per-cell rather than summed per-plate counts; and for edgeR without EB shrinkage, there are insufficient residual d.f. to stably estimate the dispersion.

**Figure 2.**
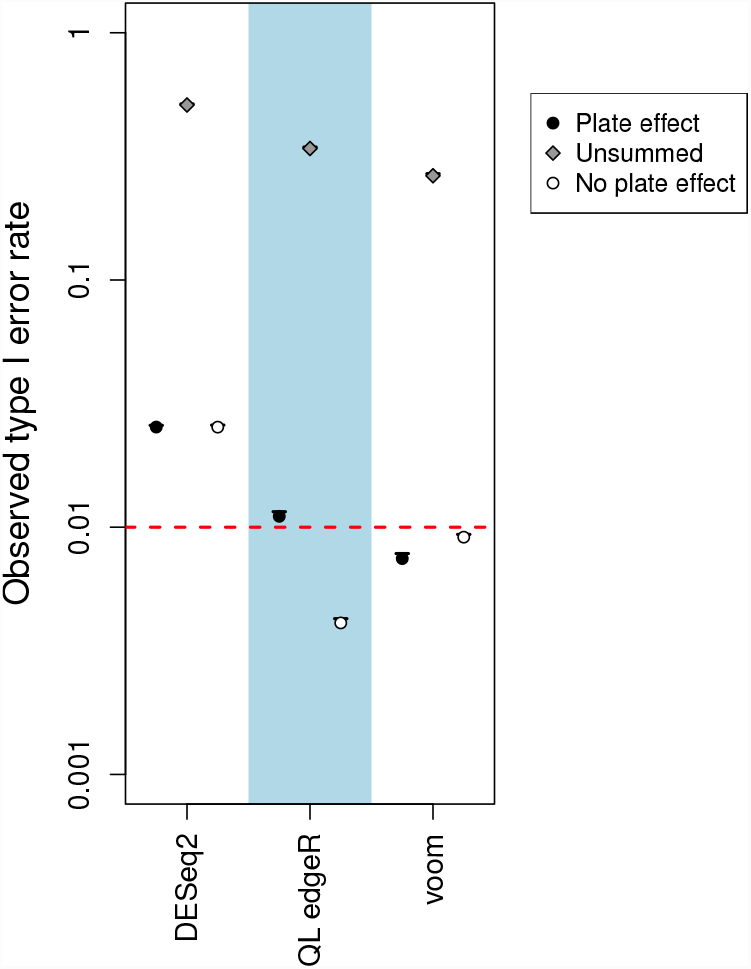
Observed type I error rate for each method after summation in simulations with and without a plate effect. Error rates are shown on a log scale and represent the average across 10 simulation iterations. Error bars represent standard errors, and the threshold of 0.01 is represented by the red dashed line. The observed type I error rate for each method without summation is also shown for comparison.

Summation will not explicitly protect against pathological situations where, e.g., *δ*_*ikg*_ > 1 for all plates in one group and *δ*_*ikg*_ < 1 for all plates in the other groups. Such genes are more likely than others to be type I errors, regardless of whether single-cell or summed counts are used in the DE analysis. However, the benefit of summation is that it provides more accurate control of these errors. This is achieved through the appropriate consideration of the model uncertainty via low residual d.f., which reduces the significance of spurious DE caused by variable *δ*_*ikg*_.

### Summation does not compromise DE detection power

It should be stressed that summation of counts does not change the nature of the underlying changes in expression. In a DE analysis, the average expression of a gene is computed for each group and then compared between two or more groups. This is true regardless of whether those groups contain replicate cells or replicate plates. In general, summation only affects the estimation of the variance rather than that of the effect size, i.e., the log-fold change.

To demonstrate, we repeated the simulations after introducing DE genes between groups. We used the various analysis methods with either the single-cell counts (i.e., ignoring the plate effect) or summed counts to detect known DE genes. For each method, the receiver-operating characteristic (ROC) curve for the analysis on summed counts was similar to that for the single-cell counts (Figure 3). This indicates that detection power at any given false positive rate is not compromised by summation. In fact, at low false positive rates, a modest increase was observed in the true positive rate of each analysis with summed counts compared to its single-cell counterpart. Similar results were observed in additional simulation scenarios involving no plate effect, stronger DE fold changes and more DE genes (Supplementary Figure S4). Moreover, the observed FDR was only controlled below the nominal 5% threshold when summation was used (Supplementary Figure S5). This is consistent with the loss of error control when plate effects are ignored.

**Figure 3.**
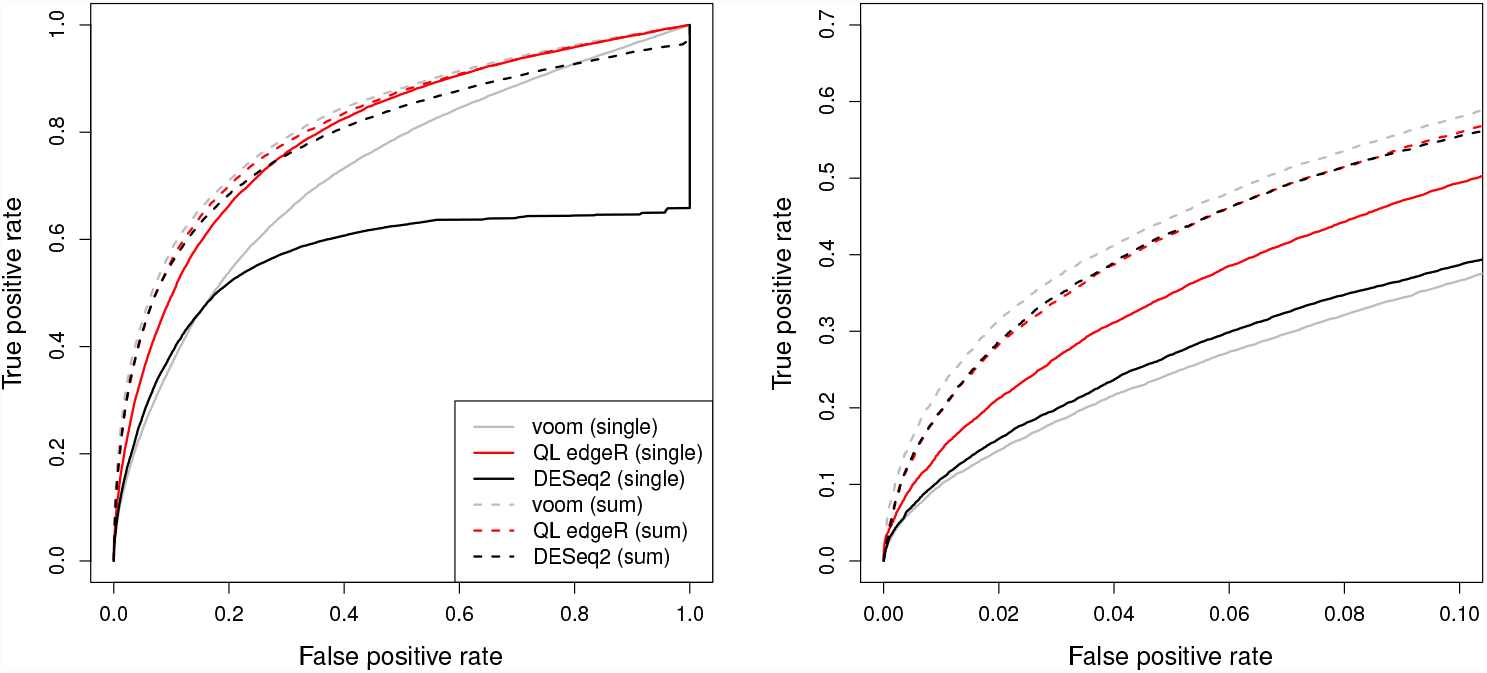
ROC curves for each analysis method on simulated data with a plate effect and genuine DE, for both single-cell (full) and summed counts (dashed). Curves are shown for DESeq2 (black), voom (grey) and QL edgeR (red). Each curve represents the average result of 10 simulation iterations. The full curves are shown on the left, and an enlarged plot focusing on low false positive rates is shown on the right.

The comparable performance of analyses with and without summation is driven by several factors. Firstly, as previously discussed, the (expected) DE log-fold change is the same regardless of whether the average expression in each group is computed over cells or over plates. Secondly, the count sum per plate is less variable than the count per cell. This is most obvious in voom where expression is quantified as a (log-)CPM value. The CPM of a count sum effectively represents an estimate of the conditional mean across all cells on a plate – by the law of large numbers, this should converge to its expectation for a large number of conditionally independent cells. Similar behaviour is observed in NB models where the variance of the sum of identical and independently NB-distributed random variables approaches that of a Poisson distribution. In both cases, the decrease in the conditional variance relative to the mean offsets the loss of power from the decrease in the number of samples when plates are considered instead of cells. (See also Section 3 of the Supplementary Materials for a discussion on how the relative decrease in the conditional variance also improves model accuracy.) Thirdly, methods like edgeR, DESeq2 and voom/limma share information between genes to estimate the dispersion or variance. This mitigates the effect of reduced residual d.f. for count sums. Finally, the increase in power at low false positive rates may be due to the greater suitability of statistical models for count sums. For example, the log-normal model in voom is more accurate at larger counts where discreteness is less pronounced.

### Justifying summation in a single-cell study

Despite the obvious improvement in performance, the summation strategy may not have unqualified appeal. After all, it seems to contradict the purpose of a scRNA-seq study. Why would one bother to sequence the transcriptome of individual cells, only to add the counts together during the analysis? The use of summed counts is equivalent to performing bulk sequencing on the pooled population, which would be technically easier to undertake and analyze. In fact, this apparent contradiction is relevant to all DE analyses of scRNA-seq data where average expression levels are compared between pre-defined groups of cells. Such averages could be obtained directly with bulk sequencing of the groups, rather than sequencing of the individual cells.

The resolution of this contradiction is based on the ability of single-cell approaches to characterize and define the groups prior to the DE analysis. Purified populations of rare cells can be profiled by scRNA-seq if they cannot be obtained in sufficient numbers for bulk sequencing. Low quality libraries or contaminating cells can be identified and removed from each group to avoid distorting downstream inferences. Groups can also be empirically identified based on gene expression patterns, though this requires some care to avoid circularity in the subsequent DE analysis. Obviously, it is not mandatory to use the summed counts exclusively. The single-cell counts can still be used for other aspects of the analysis, e.g., clustering, identifying highly variable genes. Indeed, one can exploit the cellular resolution of scRNA-seq data to characterise the nature of a DE gene detected with the summation approach – specifically, is the change in expression being driven by a particular subset of cells, or is it occurring consistently across the entire population? All of these advantages are lost when RNA sequencing is performed at the population level.

Note that summation across cells in scRNA-seq data is not without precedent. Jaitin *et al.* [5] pool single-cell expression profiles to obtain a combined vector of subpopulation-level counts for gene clustering. Klein *et al.* [7] also pool single-cell counts within each group prior to performing a DE analysis between groups, which increases the count size and stabilizes the estimates of the DE fold change. However, the use of summation to restore type I error control in the presence of plate effects has not yet been addressed in the literature.

## Summation results in reduced DE detection in real data

### Overview of the real data set and its analysis

To determine the relevance of the simulation results, DE analyses with single-cell and summed counts were performed on real scRNA-seq data from a study of mouse embryonic stem cells (mESCs) grown under three different types of culture conditions [8]. This data set contains multiple C1 chips across several batches for each culture type, where all mESCs on each chip correspond to a single culture type (serum, 2i, or a2i). Count data for all genes were obtained from http://www.ebi.ac.uk/teichmann-srv/espresso. To simplify the analysis, counts were only used from cells in the two batches where all three culture types were represented. This resulted in 60-90 cells from each of six plates (three culture types in each of two batches). To account for the batch effect, the experimental design was parameterized as an additive model with culture-specific expression terms and batch-specific blocking factors.

DE analyses were performed on the data using DESeq2, voom and QL edgeR. These methods were chosen as they could be applied on both the single-cell counts and summed counts for all cells in each plate. Prior to analysis, low-abundance genes (defined as those with an average count below 1 across all cells) were filtered out. Sample-level quality control was not applied as low-quality cells had already been removed. Each analysis method was applied with and without summation, to detect DE genes between 2i and serum or between a2i and serum. In each analysis, normalization was performed using the deconvolution method for the single-cell counts [13] or the DESeq method for the summed counts for each plate [1] – see Section 2.2 of the Supplementary Materials for more details on these normalization strategies. The set of DE genes detected at a FDR of 5% was then reported for each method.

### Numbers and ranking of DE genes are altered upon summation

In all comparisons, the number of DE genes detected with summed counts was substantially smaller than that detected with single-cell counts (Table 1). In fact, the DE list from the former was generally a subset of that from the latter. However, this does not mean that DE analyses with single-cell counts provide more power. To assess specificity, we repeated each comparison after swapping the culture labels of the samples being compared in one of the batches. Each group now contains one sample from each culture type, such that no DE genes should be present between groups. Only the methods using summation were able to control the type I error rate below the specified threshold (Supplementary Figure S6). (Some conservativeness is expected as genuine DE between culture types inflates the variances upon swapping.) In contrast, the single-cell analyses are liberal to a degree that is consistent with the simulations. This suggests that the increased numbers in Table 1 correspond to detection of false positives rather than genuine DE genes.

**Table 1.** Total number of DE genes detected by each method in each comparison at a FDR of 5%, using the single-cell or summed counts. The number of DE genes detected with both counts is also shown.

| Comparison | Counts | DESeq2 | voom | QL edgeR |
| --- | --- | --- | --- | --- |
| 2i vs. serum | Single-cell | 6243 | 9458 | 9107 |
|  | Summed | 3014 | 1710 | 946 |
|  | Both | 2488 | 1698 | 946 |
| a2i vs. serum | Single-cell | 5378 | 8850 | 8069 |
|  | Summed | 1688 | 774 | 241 |
|  | Both | 1487 | 763 | 241 |

To determine the effect of summation on the gene ranking, genes were sorted based on the *p*-values computed from each method. The identities of the top 20, 200 and 2000 genes were compared between analyses using single-cell and summed counts. In general, less than half of the top-ranking genes are shared between the two analyses for each method (Table 2). This difference is attributable to changes in variance modelling after summation, to focus on variability between plates rather than between cells. These rankings are important as the genes driving the biological differences between groups are expected to exhibit strong DE. Thus, the top-ranking genes are typically prioritized for further interpretation and investigation. Changes to the ranking indicate that summation will affect the biological conclusions that are taken from the analysis. Indeed, key pluripotency factors characterized by Kolodziejczyk *et al.* [8] are more highly ranked in analyses using summed counts compared to single-cell counts (*p* = 1.9 × 10^−4^, see Supplementary Figure S7). This suggests that summation can improve the biological relevance of DE results.

**Table 2.** Proportion of the top-ranking DE genes shared between analyses using the single-cell or summed counts. Top genes were defined as those with the smallest 20, 200, or 2000 *p*-values in each comparison.

| Comparison | Top | DESeq2 | voom | QL edgeR |
| --- | --- | --- | --- | --- |
| 2i vs. serum | 20 | 0.15 | 0.05 | 0.10 |
|  | 200 | 0.27 | 0.43 | 0.18 |
|  | 2000 | 0.49 | 0.60 | 0.48 |
| a2i vs. serum | 20 | 0.25 | 0.20 | 0.10 |
|  | 200 | 0.30 | 0.53 | 0.17 |
|  | 2000 | 0.48 | 0.60 | 0.51 |

One potential criticism of summation is that the variability between cells is hidden in the count sum. One might expect that genes with DE driven by a few outlier cells would be ranked highly, as they would not be penalized by a large cell-to-cell variance estimate. This would be inappropriate as such outlier patterns are uninteresting. However, these genes do not seem to be favoured in real data. The top set of DE genes from summation exhibit robust differences between groups (Supplementary Figure S8), whereas genes driven by outliers tend to be less significant. This is because the conditional variance of the count sum will be larger with fewer contributing cells, i.e., technical noise and heterogeneity will not average out if the sum is dominated by a few outliers. This increases the total variance and decreases the relative significance of DE for outlier-driven genes. For genes with more contributing cells, cell-to-cell variability will be hidden but this is not undesirable – see Section 4 of the Supplementary Materials for details.

## Discussion

Confounding plate effects are often present in scRNA-seq data sets but are typically ignored during the DE analysis. Our simulations indicate that this strategy is not statistically rigorous and will result in loss of type I error control. In this article, a solution is presented involving the summation of counts across all cells on each plate for each gene. The count sums can then be used in a DE analysis, effectively treating plates as individual samples. This restores type I error control and avoids the detection of excessive false positives. Summation prior to DE analyses also affects the biological conclusions of a real scRNA-seq study, by decreasing the size of the DE lists and improving the ranking of relevant genes relative to a conventional single-cell analysis.

We note that there are some situations where summation has fewer advantages. For example, cells can be arranged onto plates such that each plate contains cells from multiple biological groups. DE comparisons can then be performed directly between cells on the same plate, avoiding problems from technical variability in processing between plates. This mitigates the plate effect (though an equivalent “population effect” may be observed whereby dependencies are present between cells from the same replicate animal or cell culture) and reduces the need for summation. In experiments involving one plate in each group, summation will yield only one count sum per group. This cannot be easily analyzed by existing methods as the variance within each group cannot be modelled without replication. A direct analysis of single-cell counts is more straightforward as the variance can be estimated across cells. That said, the latter analysis is only valid under the assumption that no plate effect exists. This is because the variance estimate only accounts for variability within plates, not between plates. To verify this assumption, one would have to generate data from replicate plates such that summation would be applicable.

In general, experimental designs involving several plates nested in each group seem to provide the best compromise between statistical rigour and experimental practicality. This generates the necessary replication across populations (assuming that each plate corresponds to a replicate population) while being easier to set up than plates of mixed populations or groups. For such designs, the best way to handle plate effects is to simply increase the number of plates. This provides more residual d.f. to precisely estimate the plate-to-plate variability after summation, albeit at the cost of requiring more experimental resources. The choice of the number of plates is analogous to that of biological replicates in bulk RNA-seq experiments. This suggests that 3-4 plates per group should be sufficient in most cases. The number of cells is less important, though there should be enough cells per plate to obtain stable count sums. Summation also reduces the computational time required for the DE analysis – only a small number of plates need to be processed, instead of hundreds or thousands of cells – which may be useful in large data sets.

Summation is a simple but effective approach to overcome the problem of plate effects in a DE analysis. This complements other aspects of the data analysis that use single-cell counts, e.g., in cell clustering or to identify highly variable genes. Summation also reduces technical noise and may be a more general strategy to improve data quality when many cells are available.

## Software

All software packages except MAST are publicly available from the Bioconductor project (http://bioconductor.org). MAST is available on GitHub at https://github.com/RGLab/MAST. Simulation scripts are available at https://github/MarioniLab/PlateEffects2016.

## Supplementary Material

## Acknowledgments

The authors thank Catalina Vallejos, Antonio Scialdone and Jong Kyoung Kim for advice regarding the real data analyses and for helpful comments on the manuscript. ATLL and JCM were supported by core funding from Cancer Research UK (award no. A17197). JCM was also supported by core funding from EMBL.

